# The immunologic V-gene repertoire in mammals

**DOI:** 10.1101/002667

**Authors:** David Olivieri, Bernardo von Haeften, Christian Sánchez-Espinel, Francisco Gambón-Deza

## Abstract

From recent whole genome shotgun data of 48 mammalian species, we have used our software VgenExtractor to obtain the functional V-gene sequence repertoire in order to conduct comparative phylogenetic studies. These studies reveal a large variation in the number of V-genes across mammalian species, ranging from a mere 36 V-genes in dolphins to nearly 600 V-genes in rats. Monotremes and marsupials are the only mammals possessing an additional locus, the TRMV, apart from the seven common loci found in mammals. Also, we show evidence for the loss of the light chain loci, specifically the V*κ* chain in one microbat, and the V*λ* chain in one rodent species. Finally, we suggest different features related to the evolution of immunoglobulin and T cell receptor loci, where frequent sequence duplications are seen in the former, while preserved and undiversified lineages are observed in the latter. All the V-gene sequences described in this study are available in the public database repository `vgenerepertoire.org`.

## 1. Introduction

In the jawed vertebrate genome, specialized gene clusters exist to defend the organism against infections. Amongst them are loci containing genes that are responsible for providing the organism with the ability to recognize antigens in a specific form (Flajnik, 2002; Rothenfluh et al., 1995). This recognition process is one of the most fascinating biological phenomenon, since it evolved to recognize foreign substances, even those that do not exist in nature (Landsteiner and Jacobs, 1935). Thus, the specific immune system of jawed vertebrates can produce large variability amongst the molecules responsible for antigen recognition. From simple permutations and interchanges of a small number of genes (Hozumi and Tonegawa, 1976), specialized genetic machinery creates great diversity in the receptors of B and T lymphocytes. Indeed, this system can potentially generate billions of different immunoglobulin (Ig) and T cell receptors (TCR).

The Ig and TCR gene segments, namely the variable (V), diversity (D), and joining (J), code for proteomic structures, called variable domains that interact with antigen (Ag). Due to the limited number of V-D-J gene segments that exist, as compared with the nearly infinite number of antigens that should be recognized, these gene segments are partially reorganized in genomic processes (namely, rearrangement and somatic mutation) that modify their recognition ability (Tomlinson et al., 1996).

In most mammals, the existence of seven loci has been described or deduced. Amongst these loci, three contain V-genes for immunoglobulins (IGHV, IGLV and IGKV) (Tonegawa et al., 1 Jan. 1978) and four contain genes corresponding to the four types of TCR chains, namely TRAV, TRBV, TRGV, and TRDV (which occupies the same chromosome location as TRAV) (Davis and Bjorkman, 1988). One exception has been previously described: marsupials and monotremes possess an additional locus, called TRMV that appears to be unique to these orders (Parra et al., 2007, 2008; Wang et al., 2011).

These genes have been studied extensively to uncover the detailed mechanisms of immune recognition. The Ig V-regions recognize antigen both in soluble form and in its native configuration, while the V-regions of TCRs recognize antigen segments in a denatured form and bound to molecules of the major histocompatibility complex (MHC) (Huppa and Davis, 2003).

Antigen binding requires two V-regions whose encoding genes are not found on the same locus. Thus, the binding of an antibody to its antigen is accomplished with two V-gene products, one originating from the IGHV loci and the other from any of the light chain loci (either the IGKV or the IGLV). The same principle applies to the recognition sites of the TCR, where each domain is contributed by the V-gene from its respective locus. The TCR *α*/*β* contains two variable domains, contributed from each of the *α* and *β* chains. The same mechanism occurs within the TCR *γ*/*δ*, which contains the *γ* and *δ* chains. This genomic organization is relatively recent in evolutionary history, emerging in the first vertebrates and maintained throughout all subsequent evolutionary lineages (Rothenfluh et al., 1995).

Significant variation in the number of V-genes exists amongst species. Although the reason for this variability is unknown, a possible mechanism may be birth and death processes (Nei et al., 1997). According to this hypothesis, frequent duplications occur amongst these genes, and subsequent natural mutations produce sequence change or gene loss. From the perspective of evolution, individuals may be selected depending upon the advantages associated with these modifications. This theory is supported by observing the large number of pseudogenes that are present in these loci. These pseudogenes originated from genes that were duplicated, but subsequently were not positively selected.

Given this evolutionary hypothesis associated with the V-gene repertoire, large-scale differences may exist between species that provide indirect clues about the species’ ability to adapt to its habitat. As such, studying the number and distribution of V-genes across these loci for several species is of biological significance, since the coding products of these genes are responsible for defense against infections from a large variety of pathogens.

Identification of the complete V-gene repertoire from a sequenced genome is both difficult and tedious even within a single species (Miller et al., 1998) because there are many V-genes and they are distributed across various loci. Due to the lack of appropriate tools, such studies have been carried out in only some of these loci. We have recently developed a software tool, called VgenExtractor (Olivieri et al., 2013) that reliably and efficiently extracts V-gene sequences from sequenced genomes. With this software, it is now possible to obtain and study the V-gene repertoire from the sequence data of mammalian species available in the public domain. In this study, we have catalogued the V-gene repertoire from 48 mammals, which cover nearly the entire spectrum of the constituent orders (Figure 1). All the V-gene sequences described in this study can be obtained through a searchable database repository `vgenerepertoire.org`, which we have made freely available as a web-based application.

**Figure 1.**
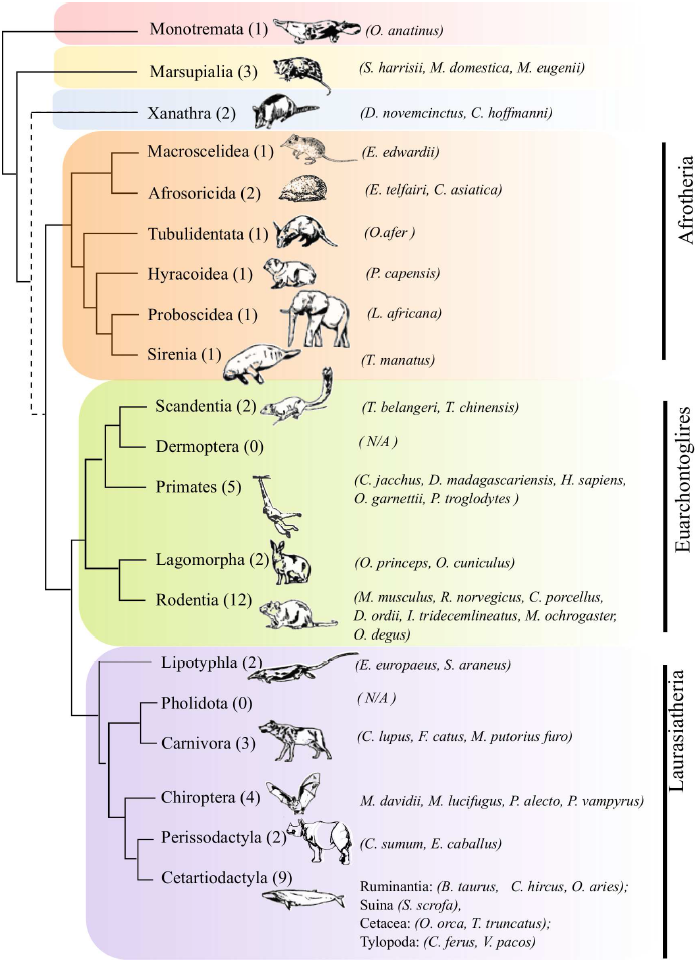
The mammalian taxa of the study. The major mammalian orders as classified by (O’Leary et al., 2013). The number of species and the precise species names that were included in this study are given in parenthesis for each of the orders. The dotted line linking the Xenarthra, indicates the present uncertainty regarding the rooting of the placental orders.

## 2. Material and methods

For these studies, we used whole genome shotgun (WGS) assemblies available at the NCBI and contigs provided in FASTA format. For most species, the complete set of V-genes from other previous studies have not been annotated, either partially or in their entirety. We used the VgenExtractor (Olivieri et al., 2013) software tool to obtain candidate V exon sequences from these large genome files. These candidate exon regions in the nucleic acid sequence are obtained by identifying RSS markers on the downstream end of the candidate exon, then searching for motifs that indicate the start/end reading frames of the V exon. To fulfill the requirement of functional V-genes, the candidate exon sequences must fulfill a set of constraints: they have an unaltered reading frame, an approximate median length of 300 bp, and contain two canonical cysteines and a tryptophan at conserved positions within the exon. During the translation of the exon region to amino acids, if a stop codon is encountered, the exon is considered non-viable and is immediately discarded, thereby avoiding the identification of pseudogenes.

In (Olivieri et al., 2013), we described the performance of our algorithm by using ground truth comparisons to publicly available data from the Ig/TCR annotation of genomes for the only two species where such annotations are believed to be complete, namely human and mouse. By assessing the false positives and negatives, we demonstrated that our algorithm detects more than 95% of the functional V regions from a sequenced genome. From these false positive/negatives, we carefully studied cases where the algorithm could fail. One source of error in the algorithm arises from the simplification applied for detecting the RSS in order to handle full genomes. In particular, from the study of six species (human, mouse, rat, anole, medaka and stickleback), the sequence of the RSS on the 3′ side of the V gene has some variations from the consensus (CACAGTG). Since most of these variations occur in the last 4 nucleotides, we locate the RSS signature from a reduced sequence (only the starting CAC sequence) (Hassanin et al., 2000; Haynes and Wu, 2004), thereby accelerating exon searches considerably. Another minor simplification pertains to canonical positions of amino acids. In particular, the algorithm constrains valid sequences to possess a canonical cysteine or tryptophan at position 41, justified by the fact that less than 1% of the six species studied lack this canonical amino acid at this position.

Apart from matching specific motif patterns along the exon sequence from whole genome scans, a fraction of the sequence extracted by the VgenExtractor algorithm may fulfill the necessary conditions described for V-genes, yet be structurally very different. Such structurally dissimilar sequences are easily discarded by using Blastp to check for sequence similarity against a known V-gene consensus, constructed from confirmed V-gene sequences from all three Ig loci and all four TCR loci. We found that even with a modest Blastp threshold (evalue = 1e-15) we could reliably identify functional V-genes.

From the functional V-genes encountered by our software, we also classified them into their respective locus type, namely, *Ig H* (IGHV), *κ* (IGKV), *λ* (IGLV) loci, TCR-*α*/*δ* (TRA/DV), *β* (TRBV), or *γ* (TRGV). Although the *α* and *δ* loci are located in the same region, we decided to treat these loci separately (locus TRAV and locus TRDV). The classification into locus type can be done using phylogenetic trees or from an automated Blastp method that searches database of known proteins. In the molecular phylogenetic approach, we aligned the sequences with clustal-omega (ClustalO) (Larkin et al., 2007), executed phyML (using LG matrix, *γ* = 2.5 and 500 bootstraps), and used MEGA5 (Tamura et al., 2011) for producing plots. The result of this analysis is that sequences from different loci naturally separate into different clades within the maximum likelihood-based tree according to their evolutionary relatedness.

These results for classifying V-genes into respective loci using phylogenetic clades have been independently corroborated with an automated software approach that we developed to classify V-gene sequences from Blastp database searches. For a given V-gene sequence, our software assigns a score for each locus based upon information from a Blastp query against the full NR protein database. The Blastp query produces a detailed listing of similar protein sequences, each consisting of a description and a similarity score. From this listing, our software calculates the probability for each locus that the V-gene sequence is of that type. This probability is based upon the total frequency of words indicative of the protein type (types of Ig or types of TCRs) found in the blastp result descriptions, weighted by the blast score for that sequence. The V-gene sequence is assigned to the locus with the highest probability. Our method is robust and consistent with the phylogenetic clade results.

## 3. Results

We have studied each of the major mammalian orders from the representative species that have been sequenced and whose data is available in the NCBI public repositories. Specifically, Figure 1 shows the species that we have studied. The classification of mammals in Figure 1 and their evolutionary origin has been taken from the conclusions of a recent study by O’Leary et al. (O’Leary et al., 2013), although we realize that the root of placentals is still disputed (Murphy et al., 2007; McCormack et al., 2012; Meredith et al., 2011; Morgan et al., 2013; Romiguier et al., 2013; Teeling and Hedges, 2013).

Together with a detailed discussion of the V-gene repertoire data obtained, we present phylogenetic trees of the V-genes for representative species of each order. For each of these trees in this article associated with the V-genes, we have indicated each locus present in mammals with different colors: red (TRAV), orange (TRDV), green (TRBV), pink (TRGV), lilac (IGHV), brown (IGKV), blue (IGLV) and gray (TRMV). In mammals the TRAV and TRDV loci are found in the same chromosomal segment, however, we treat them as different loci in the phylogenetic trees. In these trees, a group of V*δ* genes appear in the clade of the V*α*.

### 3.1. Monotremes and marsupials

The sister taxon to all remaining mammals are monotremes. Within the remaining mammals, marsupials are the closest relatives of eutherian mammals. From the order Monotremata, the platypus (*Ornithorhynchus anatinus*) genome is publicly available. With the VgenExtractor software tool, we detected 220 V-genes in the platypus (Figure 2 and Table 1).

**Figure 2.**
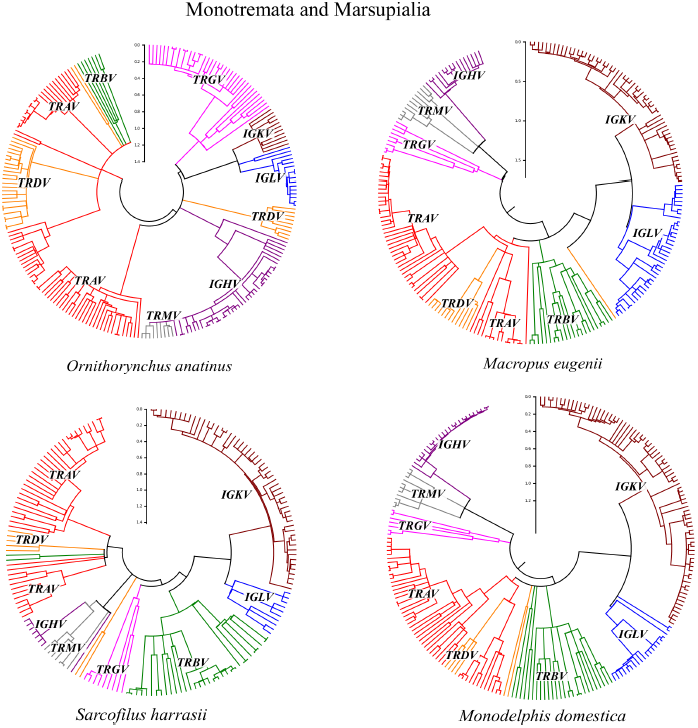
Monotremata and Marsupialia. Phylogenetic tree platypus (*Ornithorhynchus anatinus)*, wallaby (*M. eugenii*), Tasmanian devil (*S. harrisii*) and opossum (*M. domestica*) performed with the amino acid sequences obtained from the VgenExtractor program. Clades correspond to sequences found on separate loci. In mammals, the TRAV and TRDV loci are in the same chromosomal segment but are treated here as different loci. Both the monotremes and marsupials have the additional locus TRMV.

**Table 1.** Number of V-genes in one monotremes and marsupials.

| ORDER/SPECIES | IGHV | IGKV | IGLV | TRAV | TRBV | TRGV | TRDV | TRMV | ALL |
| --- | --- | --- | --- | --- | --- | --- | --- | --- | --- |
| MONOTREMATA |  |  |  |  |  |  |  |  |  |
| *O. anatinus* | 41 | 11 | 16 | 69 | 8 | 36 | 31 | 8 | 220 |
| MARSUPIALIA |  |  |  |  |  |  |  |  |  |
| *M. eugenii* | 16 | 56 | 38 | 56 | 22 | 9 | 13 | 13 | 223 |
| *S. harrisii* | 8 | 54 | 10 | 47 | 33 | 9 | 3 | 13 | 177 |
| *M. domestica* | 25 | 79 | 20 | 57 | 24 | 7 | 8 | 11 | 231 |

In the platypus, approximately 150 V-genes belong to the TCR loci. We detected only eight V-genes in the TRBV locus compared to 69 genes in the TRAV locus, representing a ratio of approximately eight to one. A group of 31 genes exists in the TRDV locus, which is a large amount compared to other mammalian species. Also, with 36 genes in the TRGV locus, the platypus has the most V-genes in this locus as compared to other mammals (Parra et al., 2006).

Given the relatively large number of genes in both the TRDV and the TRGV loci, the platypus has a large combinational potential for generating diversity in the second type of T cell receptors (the TCR-*γδ*). Also, we detected eight functional V-genes from the TRMV chain, which has been described recently in (Wang et al., 2011). From a molecular phylogenetic study, these V-genes are located within the IGHV clade, which is suggestive of their possible origin. The TRMV locus is located outside the IGHV locus. Independently, this V gene type originates from the locus of an ancient VH gene duplication. Such duplications are common and orphans V genes can be found at ectopic sites in the genomes that we studied. Some of these duplications could have been successful and generated a new functional loci, such as the TRMV, whose purpose is still presently unknown (Parra et al., 2008).

The loci of the Ig V-regions have 68 functional V-regions, where most belong to the locus of the heavy chain (41 of these VH genes have been previously studied by our group (Gambon-Deza et al., 2009)). We detected only eleven functional V-genes in the IGKV locus and 16 in that of the IGLV, in agreement with other published studies reported by Johansson et al. (Johansson et al., 2005).

In Marsupialia, there are three sequenced genomes publicly available; two correspond to the Australian marsupials (*Macropus eugenii* and *Sarcophilus harrisii)*, while the other corresponds to an American marsupial (opossum) (*Monodelphis domestica*). The functional V-gene repertoire is represented in Figure 2 and Table 1. The number of V-genes detected is approximately 220 in the Australian marsupial and 230 in the American marsupial. These three animals have a very similar genomic repertoire. In the Ig loci, the low number of genes (between 10 and 25) in the IGHV locus are noteworthy, since there is a short evolutionary distance between their members, indicating they belong to a single clade. In the three marsupials, the IGKV locus contains many more genes than the monotremes, suggesting a preferential importance of the *κ* chain.

Repertoire studies of the Ig V-genes in the *M. domestica* exists (Baker et al., 2005; Miller et al., 1998). Our study is consistent with these previous results, where one clade contains nearly all VH genes while a second clade has only one member. These previous studies only detected between 14 and 15 genes, while with our software VgenExtractor, we detected 25 genes. The discrepancy is explained by the different experimental methods: the published studies were obtained using clones from RNA, whereas we obtained our results from genomic DNA datasets. Such a discrepancy is to be expected with studies carried out with mRNA expression, since their expression depends upon the interaction with antigen. Thus, mRNA would provide only a subset of the V-genes and represent an underestimate of the actual V-gene repertoire. With respect to the V-genes from the TCR chains, the distribution of these genes is similar in the three marsupials: there are 22 to 33 sequences from the TRVB locus, 47 to 57 sequences of TRAV genes, and eight to 13 TRDV sequences, with additionally seven to nine sequences from the TRGV locus, which is similar in other mammals. Thus, the TRAV to TRBV ratio is approximately two to one. In marsupials, we detected the TRMV locus consisting of 11 to 13 V-regions, which is unique to monotremes (Wang et al., 2011) and marsupials (Parra et al., 2007).

### 3.2. Eutheria

*Xenarthra.* Xenarthra corresponds to a group of mammals that are characterized by the absence of teeth and a low metabolism. These animals originated in South America and its origin is a controversial subject associated with the rooting of the placental mammals (Murphy et al., 2007; McCormack et al., 2012; Meredith et al., 2011; Morgan et al., 2013; Romiguier et al., 2013; Teeling and Hedges, 2013; Delsuc et al., 2003a,b; Möller-Krull et al., 2007; Wildman et al., 2006; Delsuc et al., 2012). From the Xenarthra superorder of mammals, the genomes of the armadillo (*Dasypus novemcinctus*) and the sloth (*Choloepus hoffmanni)* have been sequenced and are available in the Ensembl and NCBI repositories. These genomes are assembled in scaffolds and were obtained with a slightly higher coverage of 2×. With the VgenExtractor software tool, we searched for all V-regions and used the same procedure described previously to classify each sequence into its respective locus.

No previous studies of the V-genes in this superorder exist. Figure 3 shows the phylogenetic tree and Table 2 contains the number of V-genes for these two animals. We identified 415 V-genes in the armadillo and 124 V-genes in the sloth. In both species, we found that V-genes are distributed amongst seven loci, consistent with that found in other mammals. In armadillo, the number of V-genes is very high due to the recent expansion of the immunoglobulin loci (as seen with the short branch lengths in the tree). This recent expansion is particularly evident in the IGHV and the IGKV loci, as seen in the tree with the presence of many small branches at more evolutionarily recent times. Also, there are more V-genes in the IGKV locus than in the IGLV locus.

**Figure 3.**
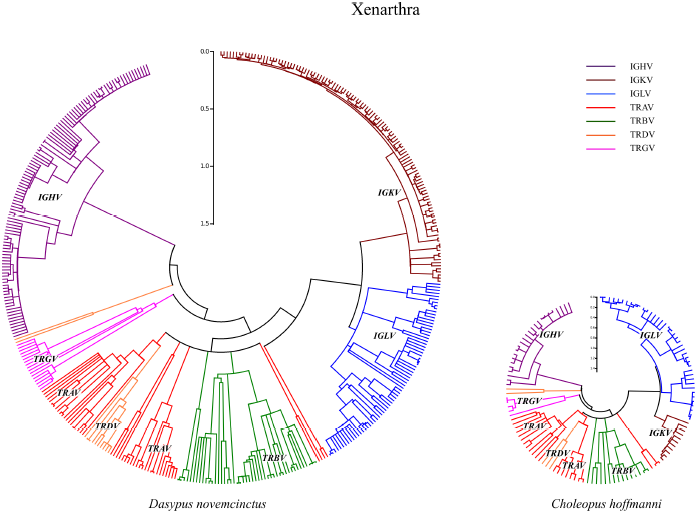
V-genes in two species of Xenarthra, the armadillo (*D. novemcinctus)* and sloth (*C. hoffmanni*). Trees were conducted with the amino acid sequences obtained by the VgenExtractor. These sequences were aligned with Clustal-omega. The phylogenetic trees were constructed from a maximum likelihood algorithm with the WAG matrix and 500 bootstrap replicates were realized for validation. Rooting was performed at the midpoint and a linearization provided in Mega5 was applied to improve the visualization of the trees. For each sequence, two independent classification techniques, described within the text, were used to determine its respective locus, thereby confirming that clades of the tree correspond to independent loci of Ig and TCR.

**Table 2.** Number of V-genes in superorder Xenarthra.

| ORDER/SPECIES | IGHV | IGKV | IGLV | TRAV | TRBV | TRGV | TRDV | TRMV | ALL |
| --- | --- | --- | --- | --- | --- | --- | --- | --- | --- |
| XENARTHRA |  |  |  |  |  |  |  |  |  |
| *D. novemcinctus* | 109 | 115 | 69 | 47 | 46 | 17 | 12 | 0 | 415 |
| *C. hoffmanni* | 25 | 11 | 40 | 25 | 15 | 5 | 3 | 0 | 124 |

Amongst the TCR loci, there are more V-genes in TRAV and TRBV than in others. Also, we found similar genes on average in the TRDV and TRGV loci in both these species, as compared with other mammals. The large difference in gene number between these two species is interesting because it may suggest different environmental evolutionary pressures.

*Afrotheria.* The Afrotheria correspond to the group of mammals that evolved in Africa. In this study we examined the V-gene repertoire from the genomes of the African elephant (*Loxodonta africana*), the Cape rock hyrax (*Procavia capensis*), the small Madagascar hedgehog (*Echinops telfairi)*, the aardvark (*Orycteropus afer)*, the Cape elephant shrew (*Elephantulus edwardii)*, the Cape golden mole (*Chrysochloris asiatica*), and the Florida manatee (*Trichechus manatus*) (Table 3 and Figure 4).

**Figure 4.**
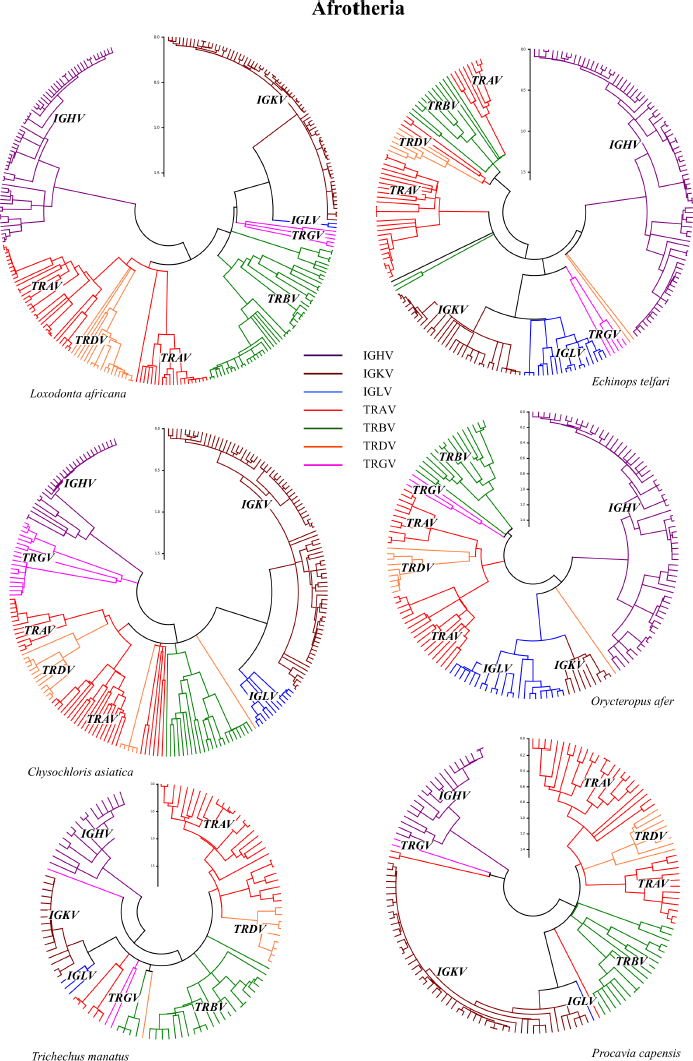
V-genes in six species of Afrotheria: (*L. africana*, *E. telfairi*, *Ch. asiatica*, *O. afer*, *T. manatus* and *P. capensis)*. Trees were constructed using the same analysis methodology mentioned previously.

**Table 3.** Number of V-genes in superorder Afrotheria.

| ORDER/SPECIES | IGHV | IGKV | IGLV | TRAV | TRBV | TRGV | TRDV | ALL |
| --- | --- | --- | --- | --- | --- | --- | --- | --- |
| AFROTHERIA |  |  |  |  |  |  |  |  |
| *L. africana* | 66 | 74 | 2 | 62 | 51 | 5 | 12 | 272 |
| *P. capensis* | 21 | 51 | 1 | 42 | 19 | 2 | 4 | 140 |
| *O. afer* | 65 | 9 | 22 | 38 | 16 | 3 | 7 | 160 |
| *E. telfairi* | 88 | 33 | 19 | 43 | 13 | 5 | 3 | 204 |
| *T. manatus* | 15 | 16 | 3 | 43 | 27 | 3 | 7 | 114 |
| *C. asiatica* | 29 | 83 | 12 | 48 | 24 | 18 | 11 | 225 |
| *E. edwardii* | 37 | 4 | 21 | 62 | 22 | 9 | 8 | 163 |

In the three species: elephant, hyrax, and manatee, the low number of V*λ* regions is noteworthy. We also found that the IGKV and/or IGHV loci are largely responsible for the variations in the total number of genes, showing that these loci may be the most versatile in processes of environmental adaptation. The manatee has very few V-genes (we found 114 genes), which is a phenomenon similar to that found in the dolphin, killer whale, and walrus, suggesting that aquatic environments of mammals could have led to a decrease in the V-gene repertoire. The V-gene repertoire in the loci of the TCR chains is similar to the average found in mammals, with the exception of an increase in the locus for the TCR *α*/*β* in the African elephant.

*Laurasiatheria.* The Laurasiatheria is a heterogeneous group of mammals that evolved in the ancient northern subcontinent of Pangea, called Laurasia in the late Triassic. Within this superorder are the Soricomorpha orders (Shrew and moles), the Erinaceomorpha (hedgehogs and relatives), the Chiroptera (bats), the Pholidota (pangolins), the Carnivora, the Perissodactyla (odd-toed ungulates), the Artiodactyla (even-toed ungulates), and the Cetacea (whales and dolphins). The first two orders, the Soricomorpha and the Erinaceomorpha, represent the oldest lineages of the Laurasiatheria. Representative genome sequences are now available for each order. Nonetheless, in these species, no previous studies for uncovering the V-gene loci exist.

In the order Soricomorpha, we analyzed the Eurasian shrew (*Sorex araneus*) (Figure 5, left), while in the Erinaceomorpha we studied the hedgehog (*Erinaceus europaeus*) (Figure 5, right). In *S. araneus*, we found 287 V-genes, where 211 genes correspond to the antibody loci, while the remaining 76 are distributed amongst the TCR loci. The expansion of antibody V-genes is distributed amongst the three loci. As with all species studied so far, these species have more genes in the TRAV locus (with 42 genes) than in TRBV (with 22 genes), with very few genes found in the TRGV (five genes) locus and the TRDV (seven genes) locus. In these orders, no V-genes were found in the TRMV locus, confirming long known results.

**Figure 5.**
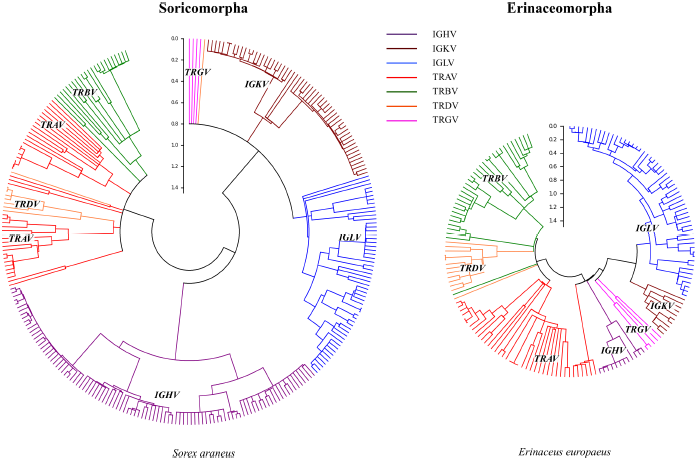
V-genes from one species Soricomorpha (*D. novemcinctus)* and one species Erinaceomorpha (*E. europaeus)*. Trees were constructed using the same analysis methodology mentioned previously.

In *E. europaeus*, we found 168 V-genes, where 77 of these genes are for Igs while the other 91 correspond to the TCR chains. Amongst the Ig genes, the number of V-genes in IGLV (55) superior to the number of IGKV (10) genes, and we detected only 12 genes from the IGHV locus. The distribution of V-gene loci within the TCR chains is very similar to the median number found in other mammals, with the highest number corresponding to TRAV compared to TRBV, and few genes in the TRGV and TRDV.

The order Chiroptera corresponds to the Laurasiatheria mammalian group that evolved the ability to fly. We studied the V-gene repertoire from two microbats (*Myotis lucifugus* and *Myotis davidii)* and two megabats (*Pteropus vampyrus* and *Pteropus alecto*) (see Figure 6).

**Figure 6.**
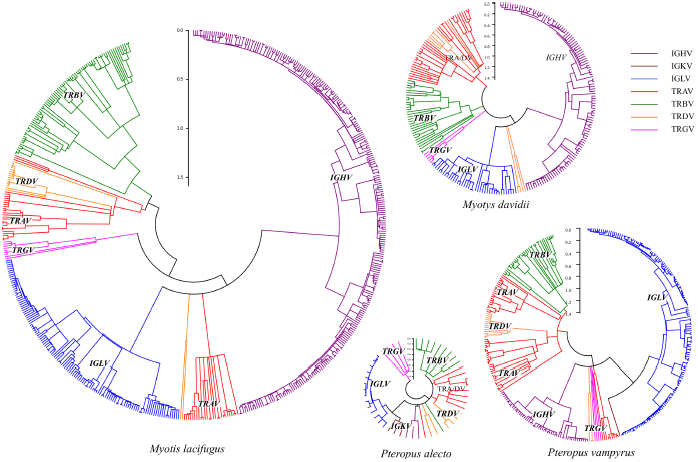
Order Chiroptera. V-genes from two species of microbats, (*M. lucifugus* and *M. davidii*), and two macrobats, (*P. alecto* and *P. vampyrus)*. Trees were constructed using the same analysis methodology mentioned previously.

Partial studies of these repertoires have been published (Bratsch et al., 2011; Baker et al., 2010). We found the presence of 554 V-genes in *M. lucifugus* (making it the mammal with the second most number of V-genes in our study, after the rat) and 234 in *M. davidii*. There is also variability in the number of V-genes between the two species of megabat: 314 in *P. vampyrus* as compared to 53 in *P. alecto*, the latter being one of the mammals with the least V-genes. Despite both being microbats, *M. lucifugus* does not have the same repertoire as *M. davidii* (Figure 6). In *M. lucifugus*, there are more V genes in all loci, with the greatest increase seen in the IGHV and IGLV loci. In the two species of megabat, *P. vampyrus* has one V-region for the *κ* chain, while in *P. alecto* we found three regions. In both cases, the viability of V*κ* genes was corroborated with the presence of the constant domain that we found from a detailed analysis of sequences in the respective genomes.

Artiodactyls and Cetaceans make up the Cetartiodactyla. Within the the Artiodactyls, we studied the sheep (*Ovis aries*), the goat (*Capra aegagrus hircus*), the pig (*Sus scrofa*), the cow (*Bos taurus*), and two camels, the Afro-Asian camel (*Camelus ferus* and the South American alpaca *Vicugna pacos*) (Figure 7). Most of the V-genes encountered In the first four species belong to TCR loci, while few V-genes were found amongst the Ig loci. In these species, there is an increase in the TCR *α*/*β* V-genes as well as genes in the TCR *γ*/*δ*. The most extreme results are found in the cow, which is one of the species that possesses the most V-genes amongst the mammals studied. It is notable that there are no V-genes in the IGHV locus, which is confirmed by other authors who have reported the low number of VH genes (Niku et al., 2012). Since our software did not detect these sequences, we carefully studied this special case in order to understand its cause. We found that the sequences have untypical alterations that manifest as changes in the canonical position 41 of the tryptophan by a cysteine, as well as possessing an unstructured RSS. As seen in Table 4, this phenomenon is only common in the Artiodactyls. The data suggest the uniqueness of the IGHV locus of Artiodactyls, which has already been described by other authors (Dufour et al., 1996). In the cow, the sheep, and the goat, the genes of the TRDV loci (and to lesser extent TRGV) have undergone a recent expansion. These results are consistent with data reported by other authors who have found the preponderance of TCR *γ*/*δ* cells in the tissue samples of bovine they studied (Mackay and Hein, 1989).

**Figure 7.**
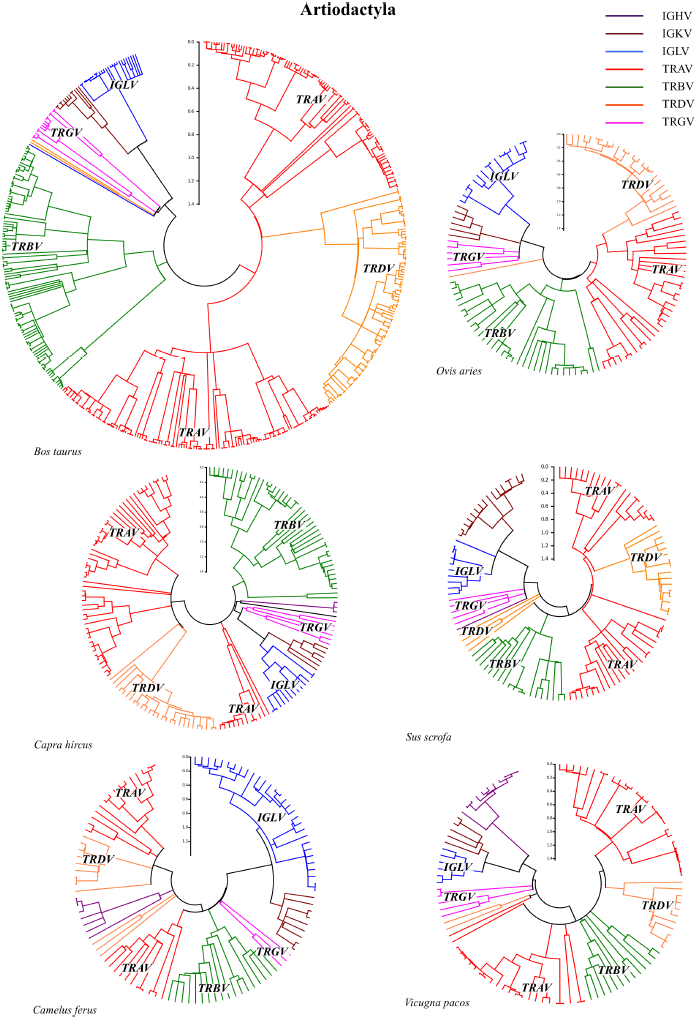
Artiodactyla. V-genes from six species: *B. taurus*, *O. aries*, *C. hircus*, *S. scrofa*, *C. ferus*, and *V. pacos*. Trees were constructed using the same analysis methodology mentioned previously.

**Table 4.** Number of V-genes in superorder Laurasiatheria.

| ORDER/SPECIES | IGHV | IGKV | IGLV | TRAV | TRBV | TRGV | TRDV | ALL |
| --- | --- | --- | --- | --- | --- | --- | --- | --- |
| SORICOMORPHA |  |  |  |  |  |  |  |  |
| *S. aureus* | 98 | 58 | 55 | 42 | 22 | 5 | 7 | 287 |
| *Erinaceomorpha* |  |  |  |  |  |  |  |  |
| *E. europaeus* | 12 | 10 | 55 | 44 | 33 | 6 | 8 | 168 |
| ARTIODACTYLA |  |  |  |  |  |  |  |  |
| *B. taurus* | 0 | 13 | 27 | 218 | 105 | 15 | 98 | 476 |
| *O. aries* | 0 | 6 | 16 | 36 | 33 | 6 | 22 | 119 |
| *C. hircus* | 3 | 6 | 13 | 63 | 45 | 6 | 28 | 164 |
| *S. scrofa* | 1 | 16 | 10 | 58 | 21 | 5 | 29 | 140 |
| *C. ferus* | 6 | 9 | 30 | 33 | 20 | 3 | 7 | 108 |
| *V. pacos* | 11 | 5 | 6 | 59 | 13 | 5 | 4 | 103 |
| PERISSODACTYLA |  |  |  |  |  |  |  |  |
| *E. caballus* | 17 | 16 | 32 | 112 | 58 | 20 | 25 | 280 |
| *C. simun* | 24 | 8 | 37 | 122 | 47 | 22 | 6 | 266 |
| CARNIVORA |  |  |  |  |  |  |  |  |
| *C. lupus* | 38 | 28 | 82 | 36 | 19 | 7 | 3 | 213 |
| *F. catus* | 24 | 13 | 47 | 46 | 20 | 5 | 7 | 162 |
| *M. putorius* | 18 | 48 | 32 | 65 | 17 | 8 | 3 | 191 |
| *O. rosmarus* | 21 | 5 | 13 | 47 | 11 | 4 | 4 | 105 |
| CHIROPTERA |  |  |  |  |  |  |  |  |
| *M. lucifugus* | 271 | 0 | 121 | 57 | 79 | 9 | 17 | 554 |
| *M. davidii* | 112 | 0 | 39 | 39 | 30 | 7 | 7 | 234 |
| *P. vampyrus* | 53 | 1 | 148 | 63 | 34 | 4 | 11 | 314 |
| *P. alecto* | 3 | 3 | 14 | 11 | 8 | 4 | 10 | 53 |
| CETACEA |  |  |  |  |  |  |  |  |
| *T. truncatus* | 2 | 2 | 10 | 10 | 9 | 1 | 1 | 35 |
| *O. orca* | 24 | 2 | 13 | 13 | 11 | 2 | 1 | 66 |

The swine group also belongs to the order Artiodactyla. Recently, the genome sequence of several pigs (*Sus scrofa*) have been uncovered and made available. With our VgenExtract software tool, we found 140 V-gene sequences. The distribution of V-genes amongst the loci is similar to that of the goat and the sheep, where a small number of Ig V-genes are found, with only one IGHV (Eguchi-Ogawa et al., 2010). In the same way, we found that the TRAV locus is greatly increased by comparison. In pigs there is a large number of TCR *γ*/*δ*, as is the case in other Artiodactyls (Uenishi et al., 2003). The two Camelidae studied have a more balanced distribution of V-genes, but the total number is low (108 in *C. ferus* and 103 in *V. pacos*). These animals have a larger number of Ig V-genes with respect to previously studied Artiodactyla, especially seen with the appearance of VH genes. In the specific case of *C. ferus*, there are more V*λ* genes than V*κ*.

Several studies (Hamers-Casterman et al., 1993; De Genst et al., 2006; Muyldermans et al., 2009; Sequencing and Consortium, 2012; Al-Swailem et al., 2010), have investigated the anomalous characteristics of Ig V domains in these species. In particular, while antibodies have been found in species of this order that are similar to those found in other mammals, additional antibody types have been found that lack the presence of the light chains in secondary immune responses (antibodies of the class IgG). However, these additional antibodies do not interfere with our results because the sequences are described by a different structure (HHV), and therefore distinct structural amino acid motifs (Hamers-Casterman et al., 1993; De Genst et al., 2006). The preponderance V-genes in the TRAV locus is maintained and we found more TCR *γ*/*δ* V-genes on average than in other species.

Cetacea are marine mammal species that evolved from Artiodactyl ancestry. Of these species, the genome sequences of the bottlenose dolphin (*Tursiops truncatus*) and killer whale (*Orcinus orca*) are available. Our analysis of these genomes uncovered only 35 V-genes in the dolphin and 66 in the killer whale (Figure 8, bottom). While both species have V-genes amongst all seven loci, they possess very few genes in each locus. Results suggest that the adaptation to aquatic environments may have influenced this decrease in V-gene abundance (Lundqvist et al., 2002) when compared with other mammals.

**Figure 8.**
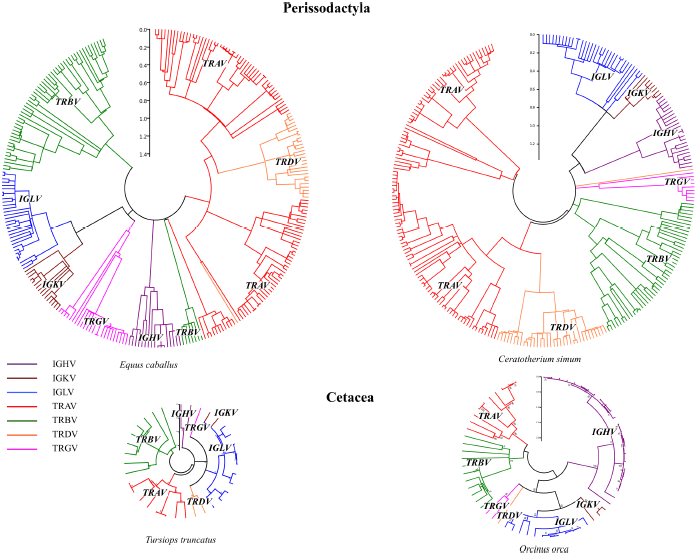
Orders Perissodactyla and Cetacea. V-genes from two species of Perissodactyla, *E. caballus* and *C. simum*, and two species of Cetacea, (*T. truncatus* and *U. orca*). Trees were constructed using the same analysis methodology mentioned previously.

The Perissodactyla corresponds to an order of Laurasiatheria. Two examples in this order are the horse (*Equus caballus*) and the white rhinoceros (*Ceratotherium simum*). We studied the genomes for each and found 280 and 266 V-genes, respectively. There is one published study from the horse where only a few IGHV genes were found (Schrenzel et al., 1997). Similar to the Artiodactyla, the number of V-genes in the Ig loci is much smaller (65 in the horse and 69 in the rhinoceros) than the number found in the TCR chains which has more than 200 V-genes. The TRBV locus increased the least, while the TRAV locus expanded the most, having more than 110 V-genes in both species. The TCR *γ*/*δ* is widely represented with 20 and 22 genes, respectively. The results in the horse are very similar to those found in Artiodactyla. Thus, both the Perissodactyla and the Artiodactyla orders possess a major variation in the distribution of V-genes: they have a low number of Ig V-genes and, in general, a large recent expansion of the TRA/DV and TRGV loci.

Carnivorans are also part of the Laurasiatheria superorder. With our software tool, we studied the V-gene distribution from the genomes of the dog (*Canis lupus familiaris*), cat (*Felis catus*), ferret (*Mustela putorius furo*), and walrus (*Odobenus rosmarus*) (Figure 9). In the antibody loci, we found that the four species have the same IGHV gene structure, where the group having the most genes must have undergone a recent expansion. In dogs, there is a detailed study of IGHV repertoire with results similar to those found in our software program (Bao et al., 2010). A surprising result is the IGLV gene expansion, especially in the dog and the cat. With respect to the TRV loci, the number of genes in TRGV and TRBV remain constant, while the size of the TRAV varies considerably, being larger in the cat, ferret, and walrus. Of the carnivorans studied, the walrus has the least number of V-genes, but maintains a similar distribution amongst the different loci compared to the rest of the carnivorans. Once again, one explanation is that the aquatic environment could have conditioned this decrease in the abundance of these genes.

**Figure 9.**
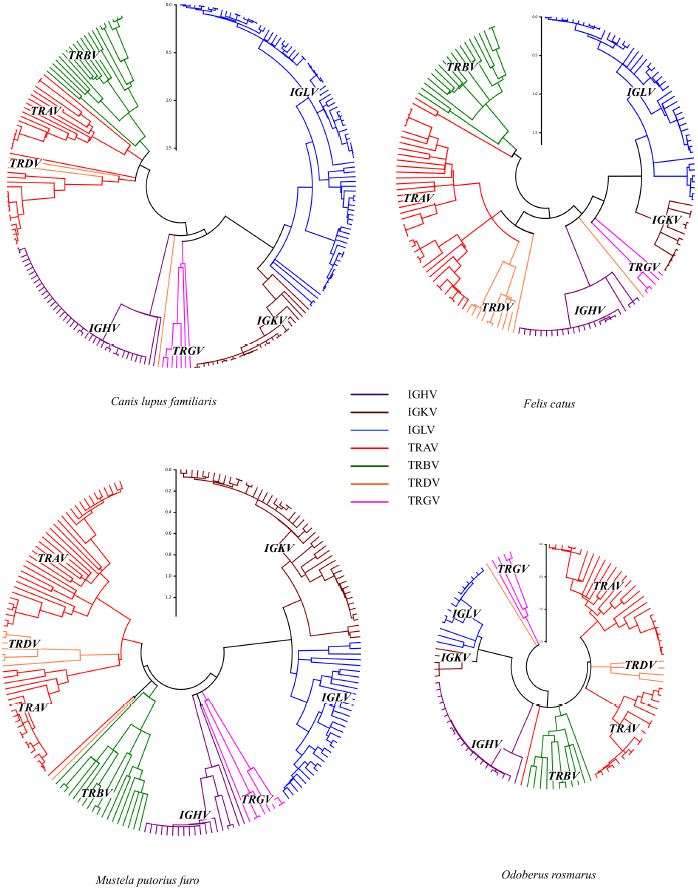
Order Carnivora. V-genes from four species: *E. Lupus familiaris*, *F. catus*, *M. putorius furo*, and *O. rosmarus*. Trees were constructed using the same analysis methodology mentioned previously.

The results indicate that in the Laurasiatheria superorder, the ancestors may have lost most genes in the IGHV and IGKV loci, and then subsequently recovered this gene loss. In this scenario, hoofed animals did not recuperate the IGHV genes and our results suggest that it may have been compensated by an increase in the V-gene repertoire of the TRV loci, notably by an increase in the number of V-genes in the *α* chain and the diversification in the second type of receptor, the TCR *γ*/*δ*. As for the carnivorans, they appeared to have recuperated from the loss of V-genes in the IGHV from frequent duplications. This hypothesis explains the fact that most genes of the IGHV locus from species within this superorder are found within a single clade. A similar mechanism must have occurred with respect to the *κ* chain (see Figure 16).

*Euarchontoglires.* The Euarchontoglires correspond to a group of mammals that evolved on the part of the Laurasia continent that is now present day Europe. Within this superorder are the Scandentia, Rodentia, Lagomorpha, Primates and Dermoptera. Within the Scandentia group, the genome sequence is available for *Tupaia belangeri* and *Tupaia chinensis*, which are small mammals that live in Southwest Asia. We analyzed these genomes with our VgenExtract software and identified 280 and 248 V-genes, respectively. Most of these genes correspond to the loci of the Ig light chains, where IGLV predominates. Between the two tupaias, variability can be seen in the number of VH genes, suggesting a recent expansion in *T. belangeri* (Figure 10).

**Figure 10.**
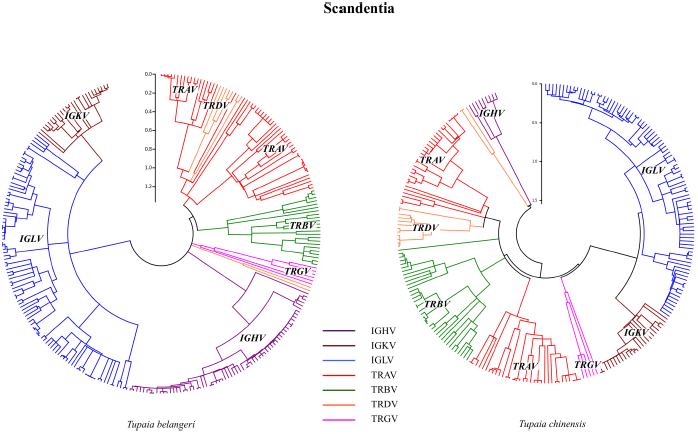
Order Scandentia. V-genes from the two species of Scandentia: *L. belangeri* and *T. chinensis*. Trees were constructed using the same analysis methodology mentioned previously.

In a previous study (Olivieri et al., 2013), we discussed the results obtained in rodents, in particular in the mouse and rat. Rodents have a large number of V-genes (Figure 11 and Figure 12); the rat has 590 V-genes, being the most numerous of the mammal that we studied. In the seven rodents studied, there exists a larger number of V-genes in the Ig loci, where all species in this order have an increase in the IGKV and IGHV loci. The IGLV locus has significant variability among the rodents. Of particular interest is the low number of V-genes from this locus found in the rat, mouse, and especially in the North American kangaroo rat *Dipodomys ordii*, where no V-genes were detected. Moreover, a further search of the *λ* chain region tested negative. Indeed, these results of *D. ordii* challenge the present belief that the Ig *λ* chains can not be lost in evolution, which was suggested from the absence of the *κ* chain in birds and snakes (Gambón-Deza et al., 2012; Wang et al., 2012). With respect to the TCR loci, our results show that there was a large and recent expansion of the TRAV locus, especially in rats, whereas in the TRBV locus, no alterations in number were detected with respect to that found in other mammals.

**Figure 11.**
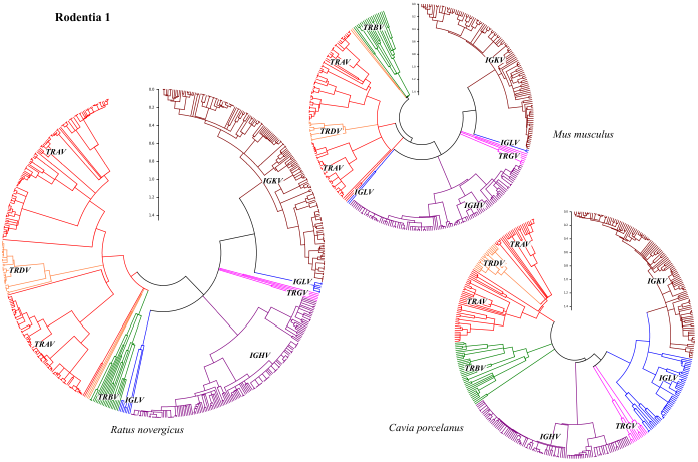
Order Rodentia. V-genes from three species of rodents: *M. musculus*, *R. norvegicus*, and *C. porcellus*. Trees were constructed using the same analysis methodology mentioned previously.

**Figure 12.**
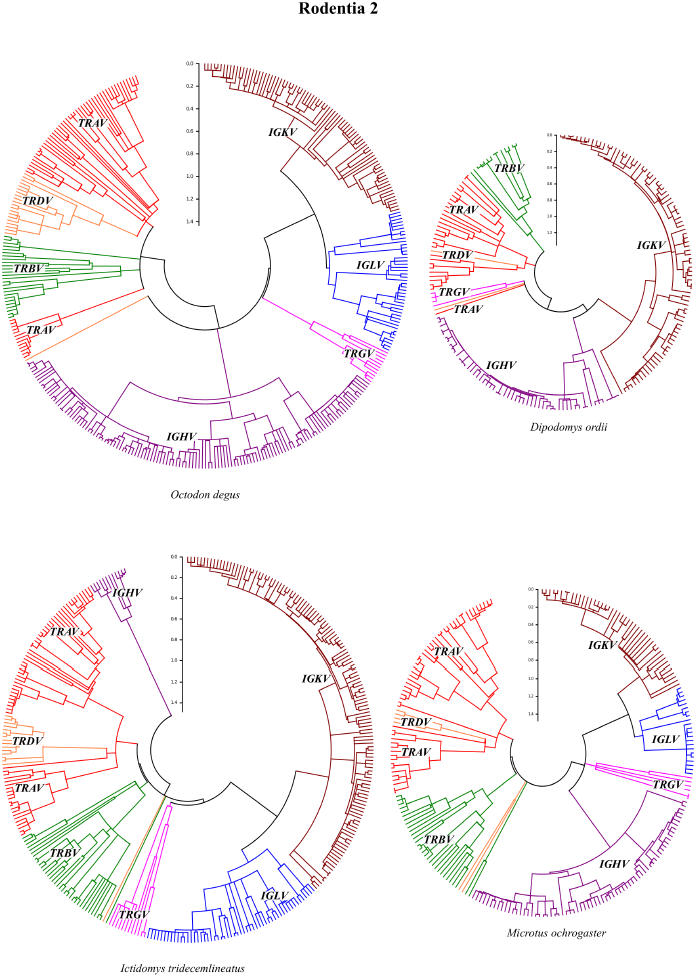
Order Rodentia. V-genes from four species of rodents: *O. degus*, *D. ordii*, *I. tridecemlineatus*, and *M. ochrogaster*. Trees were constructed using the same analysis methodology mentioned previously.

Another order of the Euarchontoglires is the Lagomorphs (Figure 13). Within this group, the genome sequence of the European rabbit (*Oryctolagus cuniculus*) and the American pika *Ochotona princeps* are publicly available. From the phylogenetic tree of the rabbit, there are a large number of V-genes in the IGHV locus. Most noteworthy is the recent and marked expansion of the three TRBV evolutionary lineages compared to other mammals. This expansion may be related to evolutionary changes in the MHC molecules previously described by Goüy de Bellocq et al. (2009). The American pika *O. princeps* has a small V-gene repertoire (102 V-genes). This animal is distinguished by having very few Ig V-genes, yet having far more V-genes for the TCR *α*/*β* (Figure 14).

**Figure 13.**
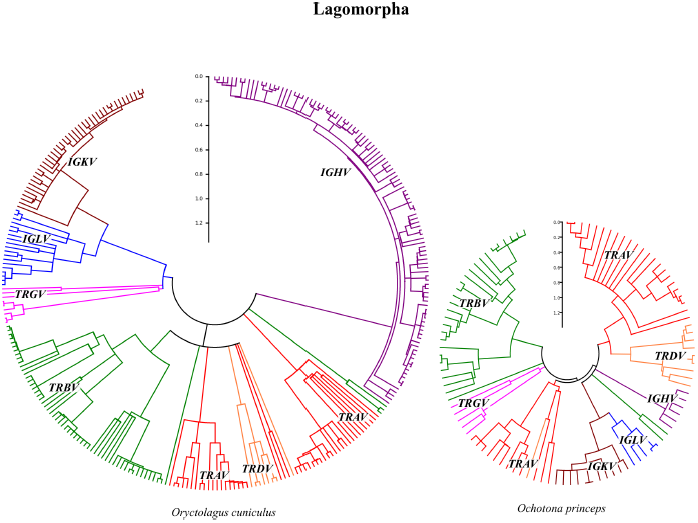
Order Lagomorpha. V-genes from two species: *O. cuniculus* and *O. princeps*. Trees were constructed using the same analysis methodology mentioned previously.

**Figure 14.**
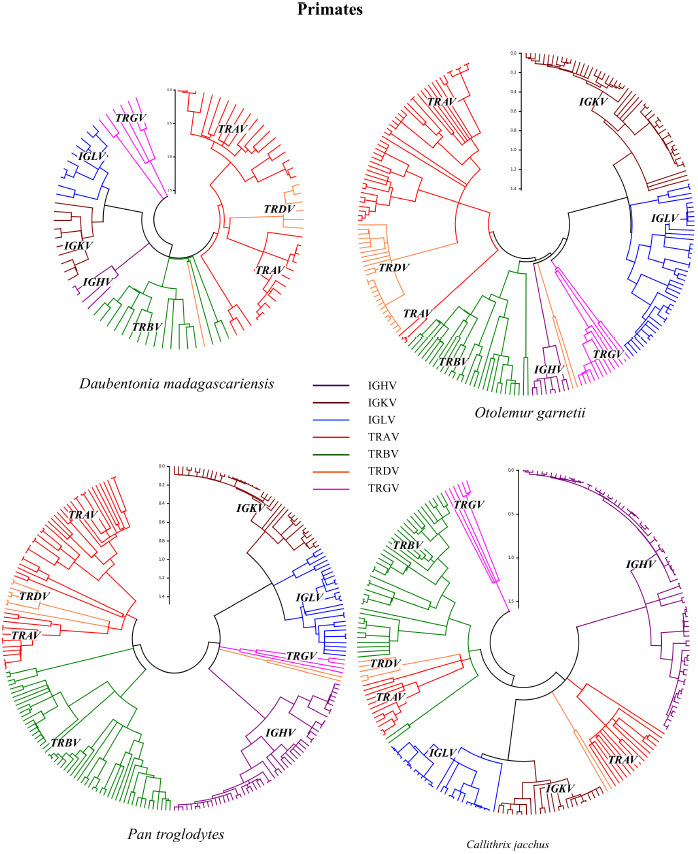
Order Primates. V-genes from four species: *O. madagascariensis*, *O. garnettii*, *P. troglodytes*, and *C. jacchus*. Trees were constructed using the same analysis methodology mentioned previously.

We studied the genome sequencing datasets from four primates: the aye-aye *Daubentonia madagascariensis*, the galago *Otolemur garnettii*, the chimpanzee *Pan troglodytes*, and the Marmoset *Callithrix jacchus*. They have approximately 200 functional V-genes evenly distributed amongst all their loci, with approximately the same number of genes in Ig and TCR loci. In (*D. madagascariensis*), the number of V-genes is half that of the other three primates studied, and while all loci are represented, most V-genes correspond to the TCR chains.

### 3.3. Evolutionary lineages of V genes

Important evolutionary phenomena between loci can be obtained from the molecular phylogenetics of V-gene sequences. The trees have distinct branch structures depending upon whether a particular locus corresponds to a TRV or IGV. The morphology of branches in trees depends upon whether or not there are frequent and recent duplications. To describe the birth and death phenomena of V-genes, we have analyzed sequence data in two ways: (a) by studying duplication events that can explain the phylogenetic morphology, and (b) by performing a multi-species analysis to show V-gene orthology.

*Duplication and expansion of loci.* Figure 15 illustrates the implication of duplication events on expansion seen in the IGHV locus of the dog (*C. lupus familiaris*) and the TRAV and TRDV loci of the cow (*B. taurus*). In this figure, genomic duplications (including both introns and exons) in the Ig and TRV loci of these species were found from an homology analysis of the genomic regions. In this analysis, homologous sequence pairs are determined by comparing a *query* sequence at a particular position with all other *target* sequences at different positions situated throughout the region. Those pairs consisting of sequences segments greater than 6000 nucleotides and having homology above 90% (but below 99.9%) are indicated in the genome regions of Figure 15 by a green track, that connects the query position to the target position. The positions of V exons are indicated in these genomic regions by red vertical lines.

**Figure 15.**
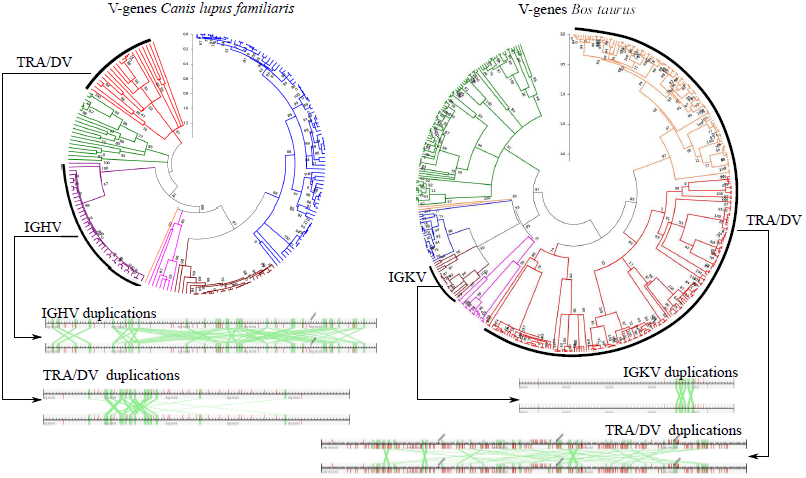
Recent Expansion of locus IGHV in dog and TRA/DV in cow. *Left:* phylogenetic tree of V-gene sequence repertoire in the dog (*C. lupus familiaris)*; the predominance of Ig V-genes is apparent where there is a recent expansion of genes in the IGHV and IGKV loci. Genomic segment duplication tracks are shown from a sequence homology studies (as explained in the text) for genomic regions of 6000 nucleotides. *Right:* a similar phylogenetic tree of V-gene sequence repertoire in the cow (*B. taurus)*. In this case, there is an expansion in the repertoire of the TCR loci. In this case, gene duplication was studied in segments consisting of 10000 nucleotides. Also, a large number of recent duplications can be seen. The IGKV locus is represented to highlight the minimal changes with respect to the TRA/DV locus.

To study the relationship between duplications and the resulting loci morphology, we compared species in the Laurasiatheria order, and found that this duplication/expansion phenomena appears to be independent of loci. Within the Ig loci, the presence of recent duplications in the IGHV and the IGKV is very significant, and is characterized by short and grouped branches (Figure 15 (left) for *C. lupus familiaris*). There is no apparent expansion in the TRA/DV locus, coincident with the observation of few duplication events. In contrast, Figure 15 (right) shows the results of *B. taurus*, where many duplication events were found in the genomic region of the TRA/DV locus, while the light chain IGKV has considerably fewer such duplications.

These results suggest differentiated evolutionary pressures for Ig and TRV loci and that the birth and death rate of V-genes is variable both with respect to loci and between orders. Moreover, the morphological patterns seen in the loci of the trees of Figure 15 are consistent with the hypothesis that birth and death processes drive evolutionary changes in these genes (Ota and Nei, 1994; Nei et al., 1997). The TRV loci are seen as long branches, many containing short tufts at their extremum, consisting of two to five genes that may have been generated from recent duplications. Such phenomena indicates that TCR V-genes may also have been subject to the same birth and death processes as those in the IGV, but more constrained over time. Thus, these branches illustrate that only minor variations are permitted, as long as they don’t violate long-standing structural restrictions enforced on the entire evolutionary lineage.

In loci with V-gene expansion and many short branches, duplication events in these loci were frequent and often included many V-genes. Gene duplication also resulted in the creation of many pseudogenes now present throughout these loci. This is observed directly in the data (see Figure 15) where several homologous duplicate pairs exist that contain a V-gene at one position, but not in the corresponding homologous sequence at the other genomic position.

*Multi-species analysis and V-gene orthology.* Throughout evolutionary history, Ig V-genes have been continuously turned over. In the case of the TRV loci, these V genes were generated at some time in the past, certain evolutionary lines were created that have endured within these genes until today. By using species having different evolutionary distances from one another, we can obtain a snapshot of V-gene orthology at different points in time. This analysis provides clues concerning the relative birth/death rates of genes within each locus and between species. This is demonstrated in Figure 16. We compared the IGHV and TRAV loci using phylogenetic trees for two primates (human and chimpanzee - having a divergence time of approximately 8 millones years ago (Ma)), two carnivores (dog and cat - having a divergence time of ∼ 50 Ma) a primate and carnivore (human and cat, a divergence of ∼ 80 Ma), where the speciation (divergence) times have been obtained from (dos Reis et al., 2012). In Figure 16 duplication processes over time have generated many paralog genes. For species chosen that are furthest apart, namely the human and cat (seen in Figure 16c), there are many orthologous genes that can be identified within the TRAV locus, while no orthologs are clearly discernable in the IGHV locus. Similarly, in a comparison between two carnivores that are more closely related, such as the dog and cat (seen in Figure 16b), once again, there is a large percentage of the genes present in the TRAV locus where orthologous genes are observed TRAV locus, while it appears that the IGHV locus is devoid of orthologs. For very closely related species, such as the human and chimpanzee (seen in Figure 16a), a large number of orthologous genes can be seen in both the TRAV and IGHV loci.

**Figure 16.**
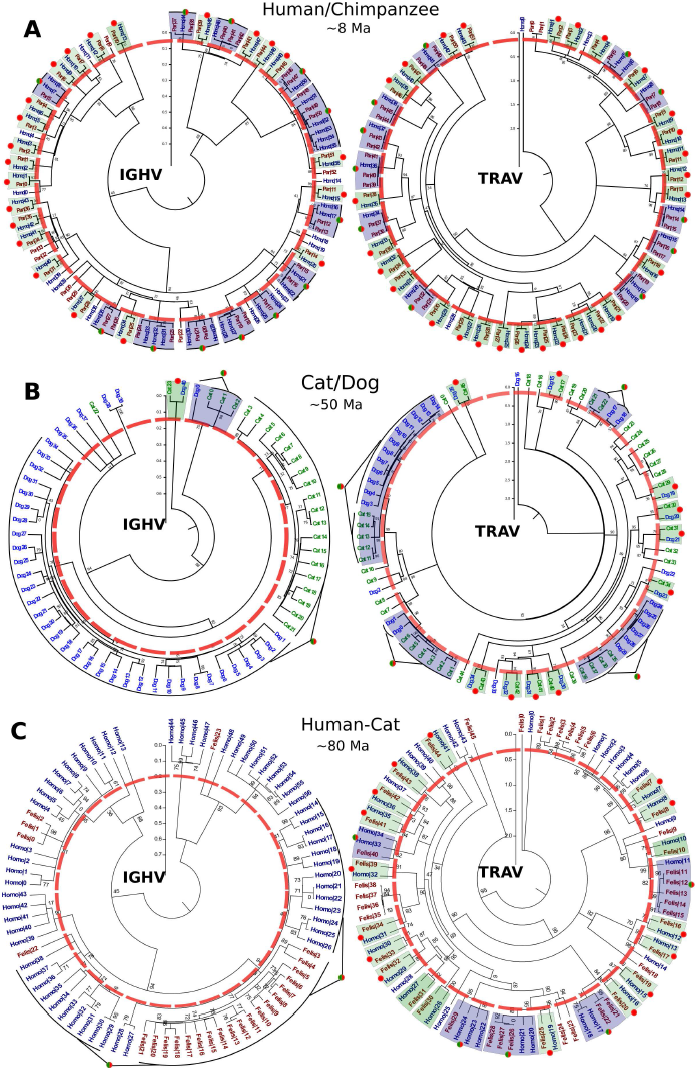
A multi-species study of the evolutionary relationships between species having different evolutionary distances from one another and orthology in the Ig and TRV loci. Phylogenetic trees of the IGHV and TRAV loci are shown for (top) chimpanzee/human, (middle) dog/cat, and (bottom) cat/human. The thick red circle with discontinuous line represents the approximate point of speciation, coinciding with the most recent divergent nodes between the two species. Orthologs have been colored (light green) and indicated by red dots, while ortholog/paralogs are colored differently (light purple) and indicated by red/green dots.

Thus, the data provided in Figure 16 shows evidence for rapid diversification of the genes in the IGHV locus with few structural restrictions, thereby explaining the rapid turnover in speciation events. In contrast, the TRAV locus are functionally constrained and must maintain certain sets of structures that allow us to identify orthologs from distant species. These multi-species representations provide a measure of the relative birth/death rates of genes within each type of loci.

The constitution of a particular evolutionary lineage in the TRV loci may represent the moment when a V-region may have started to interact with an MHC molecule. This interaction led to an evolutionary advantage that contributed to maintaining this evolutionary lineage (a co-evolution of both molecules -MHC and V-gene-). If this hypothesis is true, then it would be possible to find associations between the TCR lineages observed in the trees and different types of MHC genes. Thus, we can deduce that the orthology between evolutionary lineages of the TCR V-gene loci can be determined.

## 4. Discussion

Previous work in this area has not studied the complete genomic repertoire V-genes in mammals. Indeed, most previous studies have focused upon a single locus within a particular species. With our VgeneExtract software tool, we have been able to obtain most of these genes pertaining to the seven loci and study trends within and amongst species. There exists a variation in the total number of V-genes. The number of V-genes is one of the parameters that conditions the diversity and specificity of the immune response. In the mammals studied, some have seen an increase in the value of this parameter, larger than 300, while others have less than 100. This variation occurs even amongst species within the same order, suggesting that evolutionary pressures in each species tend to determine the number of V-genes.

Some features of the immunologic V-gene repertoire are related to concepts of the evolutionary lineage. Each species has inherited a particular V-gene group. If the sequences of this gene group had a similar modification rate as compared with all other genes within the mammalian superorders, then we should observe only small changes in the V-repertoire structure. Our results suggest that this is not the case, and that there is a preferential evolutionary mechanism which is specific to the V-genes. This can be seen by the emergence and loss of a large numbers of specific regions, such as the presence of the TRMV locus in monotremes and marsupials, the increase in the TRAV regions in Artiodactyla, the loss or underdevelopment of the IGLV *λ* chain in Afrotheria and Rodentia, or the loss of the IGKV chain locus in Chiroptera. In the case of the TRMV locus, these V-genes were created in differentiation lineage of monotremes and marsupials. This is of interest since this locus is not found in reptiles or Eutheria mammals. The most logical assumption is that the locus would be created in the common ancestor of marsupials and monotremes, which would imply a loss of this locus in the differentiation process toward Eutheria.

Regardless of the common features described, the large variability in the number of V-genes among loci is striking, even for closely related species. In the cases where loci were lost, there was always some loss of antibody light chain loci. The fact that the antigen-binding site is formed by two distinct V-genes that evolve into two independent loci have significance. The variability is greatly increased because if the antibodies are formed from a single locus they would be homodimers with two identical V-regions and therefore possess a more limited repertoire. Evolution has created different loci to increase the repertoire. In Igs, one heavy chain V domain can bind with any of the two light chains. Thus, if there are more light chain loci, then the size of the repertoire is larger.

In this study, we demonstrated that several species have lost one light chain locus (eg., *D. ordii* has lost the *λ* chain, while the *M. lucifugus* has lost the *κ* chain). Until this study, only the loss of the *κ* chains has been described in various species (the microbat, birds, and snakes), giving the false impression that there is a greater biological significance associated with *λ* chain than is the case. The results presented in this paper contradict these ideas and are the first to demonstrate species lacking the IGLV locus. In all these cases, the locus deficiency is compensated by a recent expansion of the light chain genes that remain viable or by an increased number of genes in the IGHV locus.

This work suggest a possible explanation for the differences seen in repertoire between species, namely that a species’ habitat can influence and modify the set of V-regions through mechanisms of biological evolutionary pressures. According to this hypothesis, species belonging to the same order but evolving in different environments may have very different V-gene repertoires, which are probably selected rapidly by the specific antigenic challenges of the habitat. Perhaps the clearest example that supports this hypothesis of habitat influence is provided by the genes of the rat and microbat. Both these mammals have the largest V-gene repertoire amongst the mammals that we studied. Our results suggest that a correlation may exist between the influence of the infectious environments where these animals live and the large V-gene repertoire. If this were the case, animals living in such environments may survive due to their increased number of V-genes, which contributes to generating a more protective immune response as compared with animals living in less infectious environments and with fewer V-genes. We postulate that this large V-gene repertoire has given these mammals an ecological advantage, allowing them to occupy and survive in niches that exclude other species having less immunologic protection.

Each of these species (rat or microbat) has generated a massive V-region repertoire, but within different loci. While in the microbat the IGHV locus contains the most V-genes, in the rat it is the TRAV that has the most V-genes for this species. These differences in repertoire may be explained by the types of infectious threats that these species must confront within their particular environments, since expansion of Ig is related to humoral response while an expansion of of TRV is related to cellular mechanisms. An example illustrating these ideas is seen in the *M. lucifugus*. These animals spend long periods of their lives in caves, where they hibernate gregariously and produce offspring. The septic and closed environment of these caves, tempered by the fermentation of their feces, may explain their abundant V-gene repertoire. While this interpretation is suggestive and fits the observed V-gene increase, careful studies would be needed in order to validate this hypothesis and fully understand why one species has more than 300 V-genes, while a closely related species has only 48 V-genes available for their immune system.

Similar concepts can be applied to the Artiodactyla species that have experienced an increase in the V-regions of the *γ*/*δ* receptor and a vast recent expansion of the repertoire of the TRAV locus, although we still do not know the causes in these cases.

Thus, these molecular phylogenetic studies of V-genes reveal important evolutionary clues. Throughout evolutionary history, antibody V-genes have been continuously turned over, which has conditioned the distribution of the evolutionary clans described in several studies (Kirkham et al., 1992; Anderson and Matsunaga, 1995). Such processes are not underway in the TRV loci. In genes of these loci, the birth and death mechanism occurs, but in a restricted manner. Since the product of these genes recognize antigen through interaction with MHC, evolutionary pressure may conserve parts of the sequences for this purpose.

The process of birth and death generates a large number of paralog genes in the V gene loci. Until our study, it was not possible to establish clear relationships between the Ig and TCR loci between species. Our data suggests the presence of gene orthologs and the existence of only a few paralogs in TCR loci. Because of the evolutionary implications and for the important role in generating immune repertoire diversity, these results require additional studies. With the concepts of birth and death, the elimination of a V-gene can occur without serious effects to the species since there are other concomitant duplications that maintain the potential response. In the case of a V-gene of the TRV, the situation may not be the same. Some of the evolutionary lineages appear to have only one member. The loss of this member could signify the loss of an interaction with an isotype of MHC, therefore producing a severe immune deficiency that, a priori, could not be compensated by the duplication of another line also subject to similar MHC isotype binding constraints. This implies that there are evolutionary pressures that do not permit the disappearance of the specific V-gene, which is consistent in explaining the observed lineages. Further studies should be conducted to define all lineages in the four locus of TRV. Also of interest is the evolution of these lineages and their possible co-evolution to the different MHC forms. The results given here open a new field of research concerning the structure of the repertoire of the V-genes in the evolutionary process and its relationship with the environment.

**Table 5.** Number of V-genes in superorder Euarchontoglires.

| ORDER/SPECIES | IGHV | IGKV | IGLV | TRAV | TRBV | TRGV | TRDV | ALL |
| --- | --- | --- | --- | --- | --- | --- | --- | --- |
| SCANDENTIA |  |  |  |  |  |  |  |  |
| *T. chinensis* | 9 | 25 | 90 | 71 | 38 | 5 | 10 | 248 |
| *T. belangeri* | 66 | 26 | 96 | 54 | 23 | 6 | 9 | 280 |
| RODENTIA |  |  |  |  |  |  |  |  |
| *M. musculus* | 109 | 111 | 4 | 94 | 18 | 7 | 10 | 353 |
| *R. norvegicus* | 146 | 174 | 15 | 200 | 20 | 5 | 30 | 590 |
| *C. porcellus* | 97 | 117 | 48 | 75 | 38 | 11 | 11 | 397 |
| *D. ordii* | 55 | 86 | 0 | 36 | 11 | 4 | 1 | 193 |
| *O. degus* | 125 | 80 | 44 | 62 | 26 | 16 | 12 | 365 |
| *I. tridecemlineatus* | 12 | 131 | 54 | 77 | 30 | 12 | 7 | 323 |
| *M. ochrogaster* | 69 | 46 | 21 | 61 | 31 | 6 | 3 | 237 |
| LAGOMORPHA |  |  |  |  |  |  |  |  |
| *O. cuniculus* | 108 | 42 | 19 | 51 | 59 | 9 | 4 | 292 |
| *O. princeps* | 6 | 11 | 5 | 38 | 30 | 5 | 7 | 102 |
| PRIMATES |  |  |  |  |  |  |  |  |
| *H. sapiens* | 45 | 46 | 32 | 46 | 41 | 9 | 4 | 223 |
| *D. madagascariensis* | 4 | 10 | 10 | 41 | 17 | 5 | 2 | 89 |
| *O. garnettii* | 9 | 49 | 42 | 69 | 31 | 13 | 5 | 218 |
| *P. troglodytes* | 53 | 41 | 27 | 52 | 55 | 4 | 5 | 237 |
| *C. jacchus* | 75 | 24 | 27 | 30 | 43 | 6 | 3 | 208 |

